# Bycatch mitigation could prevent strong changes in the ecological strategies of seabird communities across the globe

**DOI:** 10.1101/2021.05.24.445481

**Authors:** Cerren Richards, Robert S. C. Cooke, Diana E. Bowler, Kristina Boerder, Amanda E. Bates

## Abstract

Fisheries bycatch, the incidental mortality of non-target species, is a major threat to seabirds worldwide. Mitigating bycatch is an important factor to reduce seabird population declines and consequent changes in ocean trophic dynamics and ecosystem functioning. However, it remains an open question how and where mitigating bycatch at a global scale may conserve seabird traits and the ecological strategies that traits represent. Here we combine a dataset of species’ traits and distribution ranges for 341 seabirds with spatially resolved fishing effort data for gillnet, longline, trawl, and purse seine gears to: (1) understand spatial variation in seabird community traits; and (2) test whether mitigating fisheries bycatch may prevent shifts in traits of seabird communities and loss of ecological strategies. We find distinct spatial variation in the community weighted mean of five seabird traits (clutch size, body mass, generation length, foraging guild, and diet guild). Furthermore, our analysis suggests that successful bycatch mitigation could prevent strong shifts in the traits of seabird communities across the globe particularly in the North Atlantic and Southern Oceans. Specifically, changes in dominant foraging and diet guilds, and shifts towards communities with faster reproductive speeds (larger clutch sizes and shorter generation lengths) and smaller body masses could be avoided. Therefore, bycatch mitigation may have important indirect benefits for sustaining ecosystem functioning, as mediated by species traits. Incorporating species traits into management actions will provide valuable tools for marine spatial planning and when evaluating the success of conservation initiatives.

## 1.0 Introduction

Global fishing effort and capacity have more than doubled since 1950 (Rousseau et al., 2019) with direct and indirect ecological consequences for marine fauna (Komoroske and Lewison, 2015; Lewison et al., 2004; Senko et al., 2014). Fisheries bycatch, the incidental mortality of non-target species, is a serious threat to seabirds, (Alverson et al., 1994; Lewison et al., 2004) driving population declines worldwide (Anderson et al., 2011; Croxall et al., 2012; Dias et al., 2019; Hedd et al., 2016). For instance, the populations of three South Georgian albatross species have plummeted by 40-60 % over 35 years due to bycatch (Pardo et al., 2017). Identifying areas of high overlap between fisheries and seabirds could therefore provide critical insights for marine spatial planning and opportunities to reduce fisheries bycatch. This in turn could prevent the direct declines of seabird populations and indirect loss of ecosystem functions, such as nutrient transportation, provided by seabirds (Komoroske and Lewison, 2015).

A trait-based approach offers a valuable tool set in which to evaluate conservation successes and highlight regions where conservation strategies will provide the greatest gains. Traits are attributes of organisms measured at the individual level (Gallagher et al., 2020; Violle et al., 2007), such as body mass and foraging guild. When traits relate to function, they can be used to infer species’ contributions to ecosystem functioning (Gallagher et al., 2020). For example, seabirds are often top predators, consequently their diet and foraging strategy can relate to functions such as trophic regulation of populations and nutrient storage (Tavares et al., 2019). Thus, trait analyses may offer opportunities to highlight oceanic regions susceptible to the greatest loss of ecosystem functioning without bycatch mitigation measures.

Simple, innovative, and inexpensive mitigation solutions have substantially reduced bycatch across gear types and species (Croxall, 2008). These solutions include gear modifications that increase net visibility and deter species with scaring lines, and management actions including time-area closures that prohibit fishing in an area or at specific times (Senko et al., 2014). For example, the introduction of bird-scaring lines in a South African trawl fishery reduced albatross death rates by up to 95% (Maree et al., 2014). However, it remains an open question how and where mitigating bycatch at a global scale may conserve seabird traits and the ecological strategies that traits represent (Gallagher et al., 2020).

Here we combine a dataset of five traits across 341 seabird species with global seabird range maps and a spatially resolved fishing effort dataset for gillnet, longline, trawl, and purse seine gears to: (1) map and describe the spatial variation in community traits; and (2) test whether mitigating fisheries bycatch may prevent shifts in traits of seabird communities and loss of ecological strategies. Collectively, these objectives allow us to identify the oceanic regions potentially susceptible to the greatest loss of ecosystem functioning without bycatch mitigation measures.

## 2.0 Methods

### 2.1 Spatial data

To identify areas where fisheries and seabirds overlap, we first extracted distribution polygons for 341 seabirds from BirdLife International data zone (BirdLife International, 2017), available upon request from http://datazone.birdlife.org/species/requestdis. These spatial polygons represent the coarse distributions that species likely occupy, and are presently the best available data for the ranges of all seabirds. We subset the spatial data to only retain the extant, native, resident, breeding season and non-breeding season polygons. We created a 1° resolution global presence-absence matrix based on the seabird distribution polygons using the package ‘letsR’ and function *lets*.*presab* (Vilela and Villalobos, 2015) for further analyses. All land was removed from the presence-absence matrix using the *wrld_simpl* polygon from the package ‘maptools’ (Bivand and Lewin-Koh, 2020) and function *lets*.*pamcrop* from the package ‘letsR’ (Vilela and Villalobos, 2015).

Second, we downloaded fine scale spatio-temporal fishing effort data from Global Fishing Watch (globalfishingwatch.org). Global Fishing Watch analyses fishing activity data using the Automatic Identification System (AIS). While AIS is a safety device used onboard vessels to avoid collisions, it also transmits data about a vessel’s identity, type, location, speed and directions (Kroodsma et al., 2018). These data are processed using convolutional neural networks to characterise fishing vessel identity, gear types and periods of fishing activity with 94–97% accuracy when compared with labelled data (Guiet et al., 2019; Kroodsma et al., 2018). AIS is mandated on vessels larger than 300 gross tonnes travelling in international waters (International Maritime Organization) and is estimated to cover over 50% of nearshore and up to 80% of high sea fishing effort (Sala et al., 2018). We extracted the daily fishing activity data for gillnets, longlines, trawls, and purse seines from Global Fishing Watch. These gear types were selected because they cause the greatest seabird bycatch mortalities worldwide (Dias et al., 2019). Fishing activity between 2015 and 2018 across the four gear types was aggregated per 1° global grid cell to produce a single fishing activity layer (Fig. 1).

**Figure 1.**
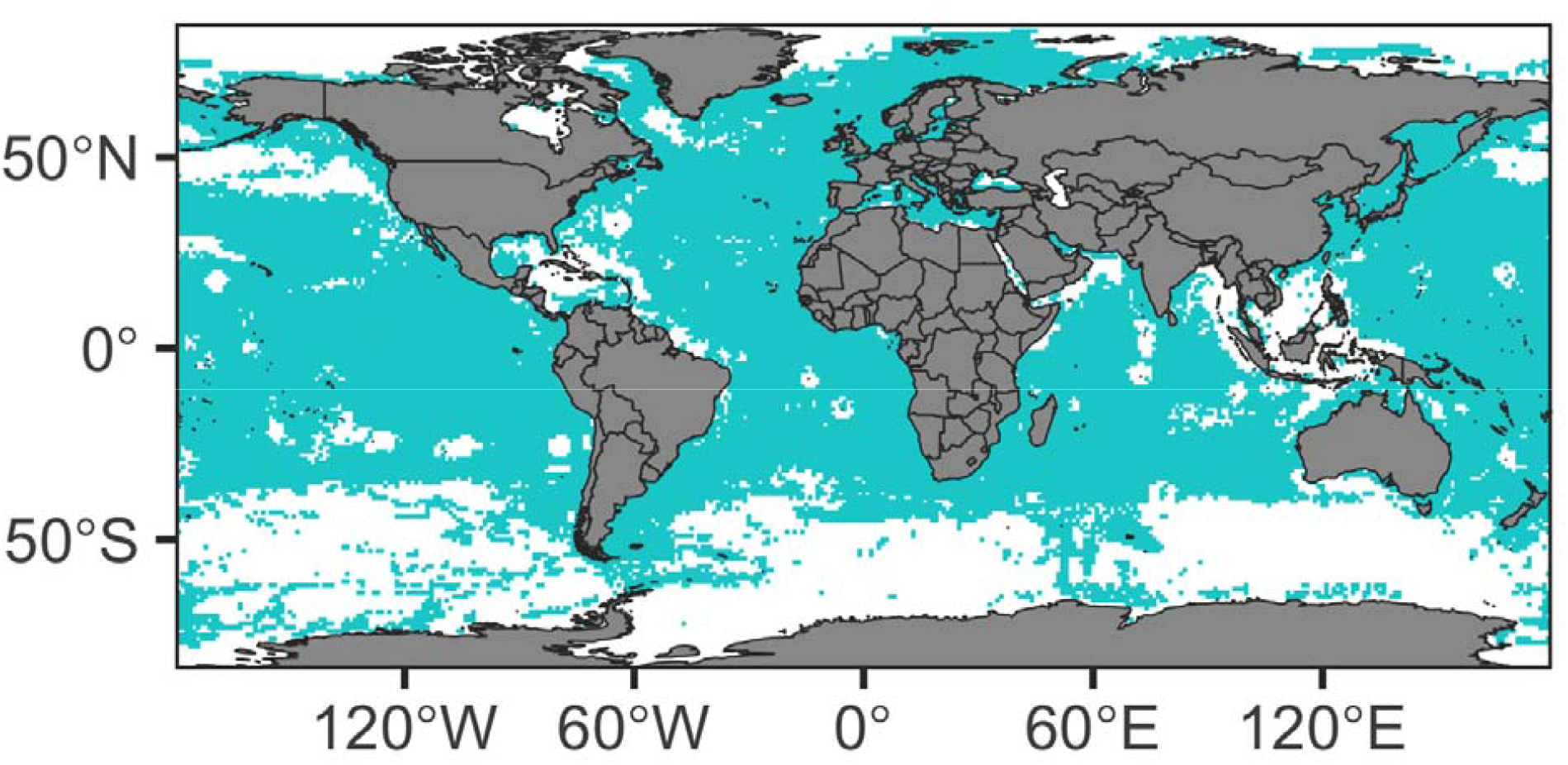
Combined distribution of fishing activity for gillnets, longlines, trawls, and purse seines identified from Automatic Identification System (AIS) by Global Fishing Watch between 2015 and 2018. Fishing effort was grouped per 1° global grid cell.

To ensure consistency between the species’ distribution and fishing activity layer, we re-projected all spatial data to a raster format with the same coordinate reference system (WGS84) and resolution (1° x 1° global grid cells). To achieve this, we used the package ‘raster’ and function *rasterFromXYZ* (Hijmans, 2020).

Finally, we built a second seabird raster representing the distributions where seabirds could become locally extinct due to bycatch. To achieve this, we removed all cells where the distributions of seabirds listed as threatened from bycatch by the International Union for Conservation of Nature (IUCN) threat classification scheme (n = 134 species, threats 5.4.3 & 5.4.4 from IUCN (2012a)) overlapped with the fishing activity layer.

### 2.2 Community Weighted Mean

To map and describe the global distribution of seabird traits, we selected five traits (Table 1): *clutch size*, the number of eggs per clutch; *body mass*, the median mass in grams; *generation length*, the age at which a species produces offspring in years; *foraging guild*, the dominant foraging strategy of the species (diver, surface feeder, ground feeder, generalist); and *diet guild*, the dominant diet of the species (omnivore, invertebrate, vertebrate & scavenger). All traits were extracted from a recently compiled and imputed dataset of seabird traits (Richards et al., 2021). Next, we calculated the community weighted mean (CWM) for each 1° grid cell with the function *functcomp*, package ‘FD’ (Laliberté & Legendre, 2010; Laliberté et al., 2014). Community weighted means describes the typical characteristics within a set of species by combining information on species’ traits and distributions (Duarte et al., 2017). For continuous data, CWM is the mean trait value of all species present in each 1º grid cell, and for categorical data, CWM is the most dominant class per trait within each 1º grid cell. We do not weight the CWM by species relative abundances because these data were not available.

**Table 1.** Description of the traits used in the present study and their relation to ecosystem functioning. Table modified from (Richards et al., 2021).

| Trait | Description | Ecosystem Function |
| --- | --- | --- |
| Clutch Size | Number of eggs per clutch | Nutrient storage |
| Body Mass | Log<sub>10</sub> (median body mass in grams) | Nutrient storage and transport |
| Generation Length | Log<sub>10</sub> (generation length in years) | Nutrient storage |
| Foraging Guild | The dominant foraging guild of the species<br>*Diver; Surface; Ground; Generalist* | Nutrient storage; Trophic-dynamic regulations of populations |
| Diet Guild | The dominant diet of the species<br>*Omnivore; Invertebrate; Vertebrate & scavenger* | Nutrient storage; Trophic-dynamic regulations of populations |

### 2.3 Trait shifts

To quantify the extent mitigating fisheries bycatch could prevent shifts in seabird traits, we calculated the deviation between the observed CWM (i.e., all species) and CWM after the removal of bycatch-threatened seabirds from cells overlapping with fishing activities, for each 1º grid cell. For continuous traits (clutch size, body mass, generation length), the percentage deviation in CWM for each grid cell CWM (Deviation_Continuous_) was calculated with the following equation:

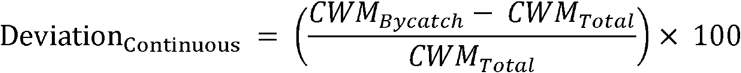

Where *CWM*_*Bycatch*_ is the community weighted mean after removing bycatch-threatened seabirds if the cell overlapped with fishing activities, and *CWM*_*Total*_ is the observed community weighted mean.

To quantify the shift in categorical traits, we calculated the proportion of each foraging (*diver, surface, ground, generalist)* and diet guild (*omnivore, invertebrate, vertebrate & scavenger)* category per grid cell as observed (Proportion_Total_), and again after the removal of bycatch-threatened species for cells overlapping with fishing activities (Proportion_Bycatch_). We then calculated the deviation between these values per grid cell (Deviation_Catagorical_) with the following equation:

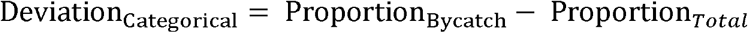

To describe the spatial trends in trait shifts across latitude, we fitted either general additive models (GAM) using the package ‘mgcv’ and function *gam* (Wood, 2017), or additive quantile regression models using the package ‘qgam’ and function *qgam* (Fasiolo et al., 2017) depending on the most appropriate fit. For each trait, latitude was included as the predictor and Deviation_Continuous_ or Deviation_Categorical_ as the response. All analyses were completed in R version 3.5.0 (R Core Team, 2020).

## 3.0 Results

### 3.1 Spatial variation in community traits

We find large spatial variation in the community weighted mean (CWM) of clutch size, body mass, generation length, foraging guild and diet guild traits across the globe (Fig. 2). Species with the largest clutch sizes are distributed along coastlines, particularly in the Northern Hemisphere. In contrast, species with the smallest clutch sizes are highly pelagic and distributed across all oceans. The CWM for body mass is more evenly distributed, with the heaviest species being located in the Southern Ocean. Species with small body masses are distributed between 30°N and 30°S. Generation length is also evenly distributed globally. Species with the longest generation lengths are concentrated in the Southern Ocean, whilst the shortest generation lengths are along coastlines. For the foraging guild, surface foragers typically dominate most oceans below 50°N, whilst divers are the most dominant above 50°N and along the coast of Atlantic Central America and Oceania. Generalists are concentrated around the coasts of Europe (Mediterranean Sea, Black Sea, Baltic Sea, North Sea) and ground foragers dominate in the high Arctic. For the diet guild, vertebrate & scavenger consumers dominate the Southern and Pacific Oceans, while invertebrate consumers dominate the Atlantic and Indian Oceans. Omnivores are only dominant in the Caspian Sea.

**Figure 2.**
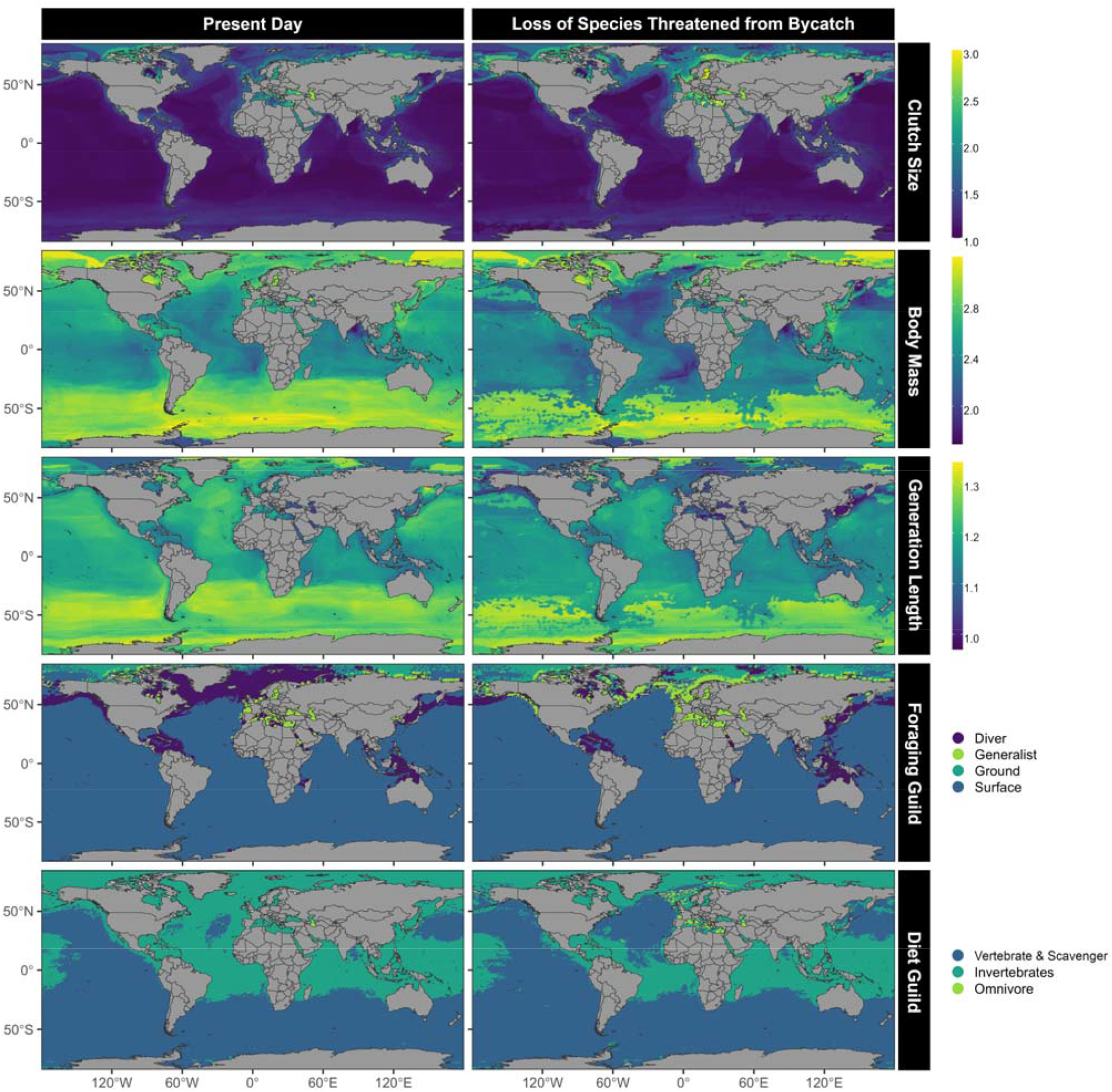
Community weighted mean (CWM) of five traits based on the distributions of 341 presently extant seabird species (left panels), and following the predicted local loss of 134 species threatened from bycatch in areas where their distributions overlap with fishing activity (right panels). Therefore, the difference represents the shifts in traits that may be prevented through successfully mitigating seabird bycatch. For continuous data, CWM is the mean trait value of all species present in each 1º grid cell. For categorical data, CWM is the most dominant class per trait within each 1º grid cell. Body mass and generation length traits are log_10_ transformed, and truncated to the 0.1 - 99.9% range to aid visual clarity. Maps including all the data are in Appendix 1, Fig. S1.

### 3.2 Preventing trait shifts

Our results suggest that successful bycatch mitigation has the potential to prevent shifts in the traits of seabird communities across the globe (Figs. 2 & 3). Removal of bycatch threats could prevent an increase in clutch size above 50°S, and a decrease in clutch size below 50°S (Fig. 3A). Furthermore, the global shift in CWM to species with shorter generation lengths and smaller body masses could be avoided (Figs. 3B-C). This trend is particularly prominent between 30° and 70° in both hemispheres. Bycatch mitigation could further prevent shifts in diet guilds from a dominance of invertebrate consumers to vertebrate & scavenger consumers in the North Atlantic Ocean, and shifts to more omnivores around European waters (Figs. 2 & 3D-F). For foraging guild, shifts from a dominance of diving and surface foragers to generalist and ground foragers above 40°N could be prevented (Figs. 2 & 3G-J). Below 40°N, we find surface foragers may remain the dominant foraging guild without bycatch intervention (Fig. 2), but their proportion within the community could increase (Fig. 3G). The proportion of divers within the community may also increase slightly between 40°N and 50°S, but decrease below 50°S (Fig. 3H). Generalist foragers could decrease below 40°N (Fig. 3I), and ground foragers may remain relatively stable (Fig. 3J).

**Figure 3.**
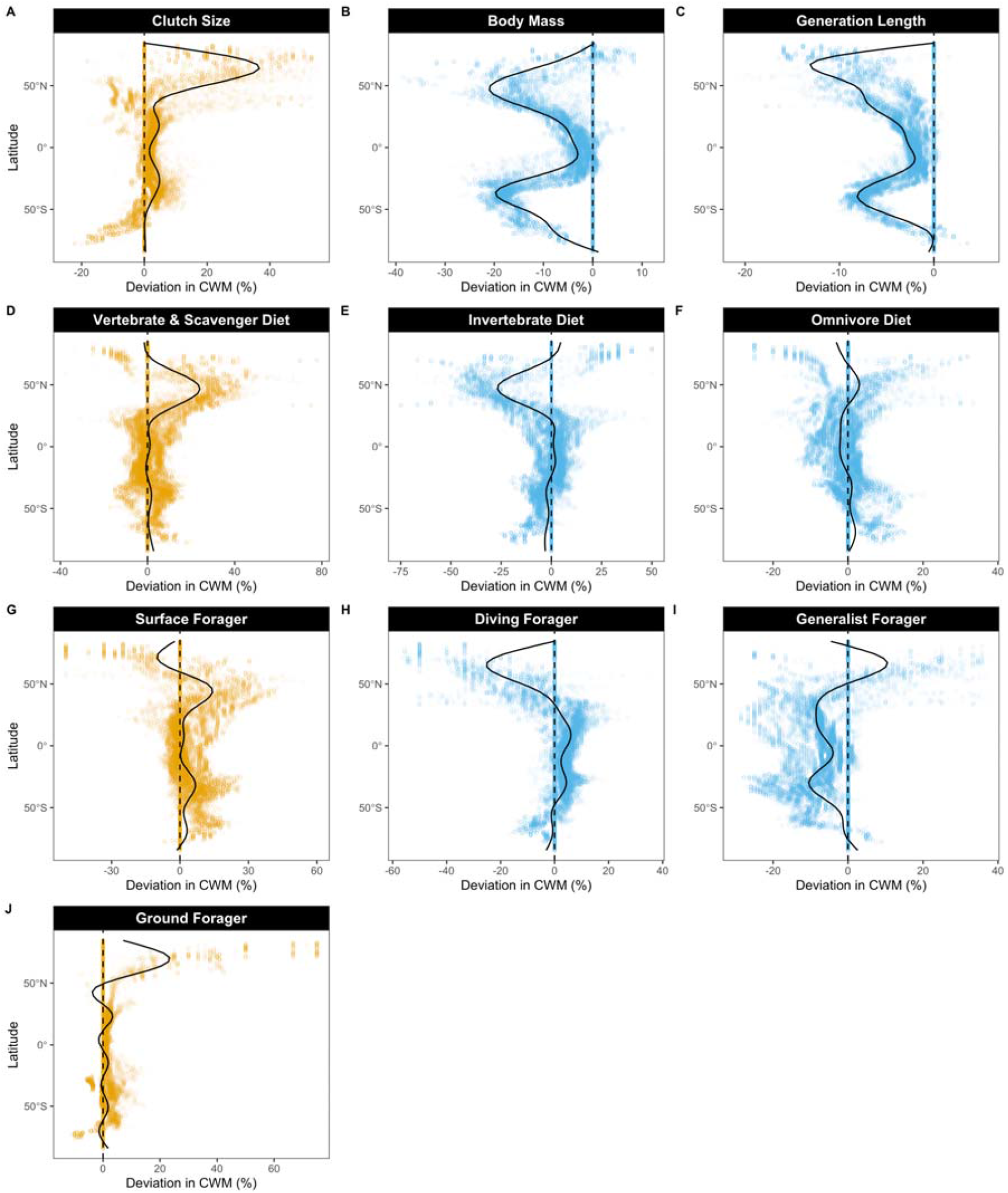
Shift in community weighted mean (CWM) across latitude following removal of 134 species threatened from bycatch in areas where their distributions overlap with fishing activity. Each data point represents the CWM within a 1º grid cell. Dashed zero line represents the CWM of the total species list (341 species). Solid black lines in A and D-J are fitted generalized additive models, and in B-C are additive quantile regression models describing the spatial trends in trait shifts across latitude. Orange represents a significant overall positive shift from the model output, and blue a significant overall negative shift in the CWM following removal of species threatened from bycatch. Figures were truncated to the 0.1 - 99.9% range to remove extreme values. Figure which include all values are in Appendix 1, Fig. S2.

## 4.0 Discussion

Here we find that mitigating bycatch could prevent large shifts in traits of seabird communities. Specifically, changes in dominant foraging and diet guilds, and shifts towards communities with faster reproductive speeds (larger clutch sizes and shorter generation lengths) and smaller body masses could be avoided. Therefore, bycatch mitigation may have important indirect benefits for sustaining ecosystem functioning, as mediated by species traits. For example, body mass is strongly linked to nutrient transport and storage because large individuals hold and disperse large nutrient quantities (Anderson et al., 2011; Doughty et al., 2016; Tavares et al., 2019). Consequently, preventing shifts to smaller body masses could protect important zoogeochemical cycles of major elements worldwide (Speakman, 2005; Wing et al., 2014; Graham et al., 2018; Schmitz et al., 2018; Tavares et al., 2019). Moreover, preventing shifts to species with faster reproductive speeds (decreased generation lengths and increased clutch size), such as gulls and tern, could have implications for nutrient storage and cycling, and food provisioning (Tavares et al., 2019). Additionally, the conservation of foraging strategy and diet guild traits may sustain trophic regulations and community structures, because, as top predators, seabirds influence marine food webs from the top down via direct and indirect pathways (Ripple et al., 2017).

Few management actions have incorporated biological trait analyses for marine spatial planning and when evaluating the success of conservation initiatives (Miatta et al., 2021). Moreover, a number of studies highlight the importance of including biological communities and ecosystem functions into conservation policy because focusing on biodiversity metrics alone may exclude functionally and ecologically important locations (Bremner et al., 2006; Frid et al., 2008; Miatta et al., 2021; Rees et al., 2012). Thus, considering the traits of species assemblages in a quantitative framework offers valuable tools for advancing marine conservation outcomes (Miatta et al., 2021). While trait-based approaches do not directly quantify the ecosystem functions that seabirds deliver, here we show biological traits could provide new insights for assessing the success of bycatch mitigation simply because fishing non-randomly targets species with different traits (Richards et al., 2021; Zhou et al., 2019). Our findings identify the oceanic regions potentially the most susceptible to shifts in seabird ecological strategies and ecosystem changes without improved conservation measures. We find the North Atlantic Ocean may be particularly vulnerable to shifts in all five seabird traits, and the Southern Ocean may be at risk to changes in reproductive speed and body mass traits without bycatch mitigation methods. These finding could be incorporated into management actions to inform marine protected area design with the intent to protect the ecosystem functions which seabirds provide.

Our results further provide a basis to advise conservation interventions, such as actions that align with the IUCN Conservation Actions Classification Scheme (IUCN, 2012b), which presents a hierarchical structure of conservation actions presently needed for species. For instance, through *water protection* actions (IUCN Action 1), the locations of greatest ecological strategy shifts could help guide where to designate and prioritise Marine Protected Areas (MPAs), therefore reducing seabird conflict with fishing gears. Our findings could further inform *water management* actions (IUCN Action 2) through identifying areas which would benefit from bycatch mitigation methods, such as fishing gear modifications, and fisheries closures during sensitive times for seabirds. Future studies and management actions may consider assessing the response of trait and ecosystem function shifts to bycatch at the local scales in these regions. For example, large-scale closure of the eastern Canadian gillnet fishery saw a positive population response in a number of seabird species (Regular et al., 2013). This example could provide a valuable case study to quantify the response of conserving whole species assemblage for regional ecosystem functioning.

We focus solely on fishing threats because bycatch is named the greatest threat to seabirds worldwide, however, seabirds face a diversity of threats throughout their life including invasive species, climate change, and pollution (Dias et al., 2019). Future studies may consider investigating how managing threats through space and time conserves seabird traits and ecosystem functions. For example, coupling extensive seabird tracking data with colony-specific trait information and regional threat patterns could provide a powerful and informative tool for local management. Finally, since our approach assumed the complete removal of species which are threatened from bycatch in areas overlapping with fishing activities, i.e. local extinction of these species, future studies may consider investigating how reduced population sizes and changes in proportions of species abundance caused by bycatch could influence community traits.

The Global Fishing Watch layers provides unprecedented understanding of the global fishing fleet and its spatiotemporal variations (Kroodsma et al., 2018). Consequently, the dataset is an invaluable resource to advance our understanding of fisheries bycatch on seabirds and other marine organisms. For example, here through integrating these fine-scale fisheries data with seabird traits and distribution data, we provided a new perspective on the potential successes of bycatch mitigation for conserving species ecological strategies and opportunities to identify oceanic areas of conservation priorities. Similarly, recent research employed the Global Fishing Watch data and biologging data to detect albatross association and encounters with commercial fishing vessels in the North Pacific Ocean, thus further revealing the fishing dataset’s value as a novel conservation and management tool (Orben et al., 2021). We encourage use of fine-scale spatial datasets to provide additional angles for seabird research, and to expand on the present study. For example, fishing activity and seabird distributions vary at different time scales, with distinct diurnal, seasonal, and annual patterns. Incorporating finer-scale data (e.g., biologging data) which encompasses these temporal signals is a direction for future studies that may provide further insights into the impacts of fishing on changes to seabird ecological strategies. Moreover, we focus on the combined distribution of gillnet, longline, trawl, and purse seine fishing activity on the overall shift in seabird traits. Additional research may consider quantifying the response of seabird ecological strategies to the spatiotemporal variations in individual gear types and intensities. Finally, the Global Fishing Watch data are fundamentally constrained by the limitations of AIS including incomplete satellite coverage in some regions, device tampering, and not all vessels carry an AIS transponder. Additionally, distributions of small-scale subsistence, and illegal, unreported, and unregulated (IUU) fishing activities were unavailable, and therefore not included in our study. However, as the AIS datasets are improved with time and fine-scale IUU fishing data become available, new patterns of seabird ecological strategy changes could be revealed.

Overall, we use extinction simulations to infer how a trait-based approach has potential to provide a unique perspective on the success of bycatch mitigation across the global seabird species pool. Specifically, we find traits can be used to identify regions potentially at risk to shifts in the ecological strategies of the species which dominate, and the impact of these species on ecosystem functions without bycatch mitigation strategies. Management actions that incorporate species’ traits and fine-scale fisheries datasets as tools for marine spatial planning will add an important dimension when evaluating the success of conservation initiatives.

## Data Sharing and Accessibility

Seabird traits were extracted from Richards et al. (2021), specifically https://doi.org/10.5061/dryad.x69p8czhd. Species distribution polygons are available upon request from http://datazone.birdlife.org/species/requestdis. Fishing effort data for 2015 and 2016 are available for download, and data for 2017 and 2018 are available upon request from https://globalfishingwatch.org/. Please contact Cerren Richards for access to R code.

## Notes

### Competing Interest Statement

The authors have declared no competing interest.

