## Appendix 1 for "Bycatch mitigation could prevent strong changes in the ecological strategies of seabird communities across the globe"

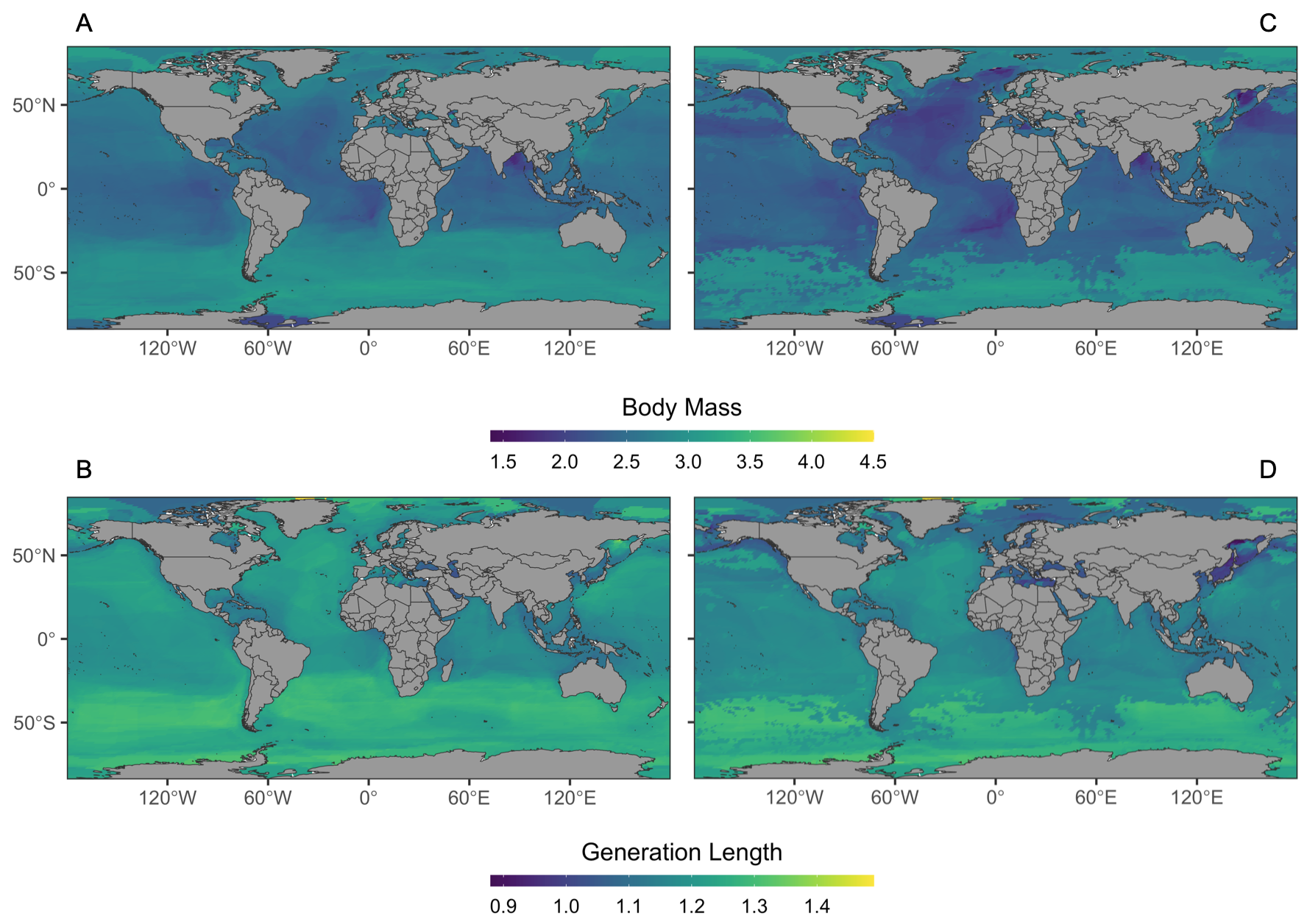

**Figure S1 |**	Untruncated community weighted mean of body mass (A & C) and generation length (B & D) traits based on the distributions of 341 extant seabird species (A & B), and following the predicted loss of 134 species threatened from bycatch in areas where their distributions overlap with fishing activity (C & D).

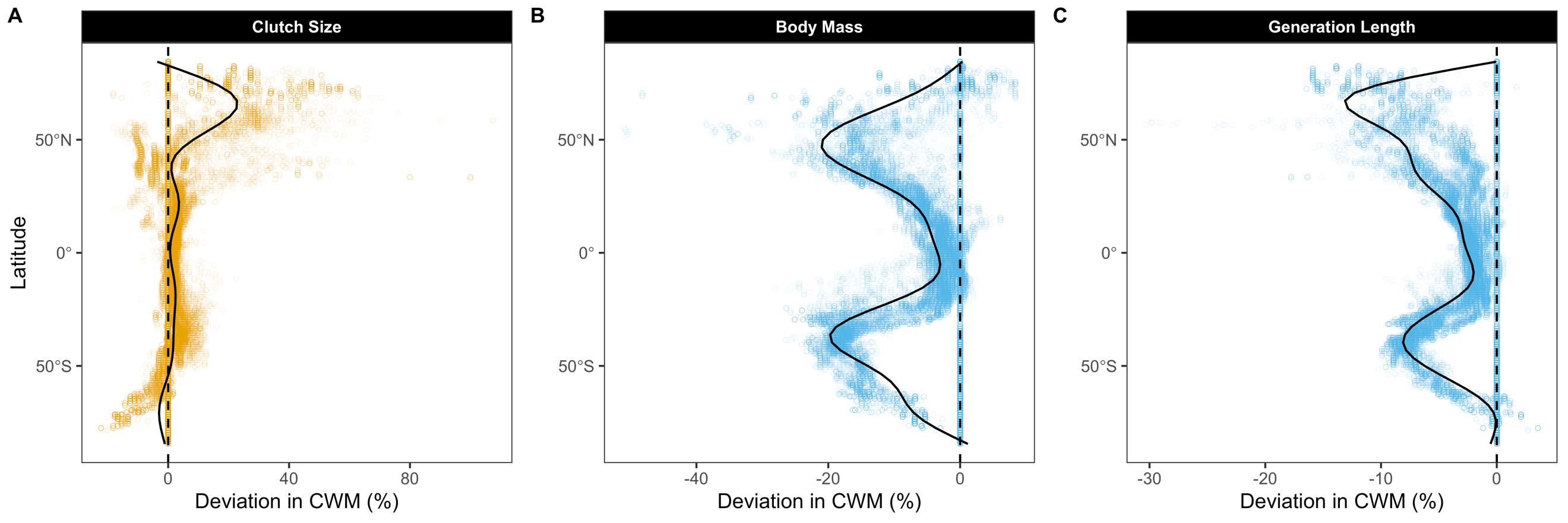

**Figure S2 |**	Untruncated shift in the community weighted mean of clutch size (A), body mass (B), and generation length (C) across latitude following removal of 134 species threatened from bycatch in areas where their distributions overlap with fishing activity.
